# Permissive zones for the centromere-binding protein ParB on the *Caulobacter crescentus* chromosome

**DOI:** 10.1101/172932

**Authors:** Ngat T. Tran, Clare E. Stevenson, Nicolle F. Som, Anyarat Thanapipatsiri, Adam S. B. Jalal, Tung B. K. Le

**Affiliations:** Department of Molecular Microbiology, John Innes Centre, Norwich, NR4 7UH, United Kingdom; Department of Biological Chemistry, John Innes Centre, Norwich, NR4 7UH, United Kingdom

## Abstract

Proper chromosome segregation is essential in all living organisms. In *Caulobacter crescentus*, the ParA-ParB-*parS* system is required for proper chromosome segregation and cell viability. The bacterial centromere-like *parS* DNA locus is the first to be segregated following chromosome replication. *parS* is bound by ParB protein, which in turn interacts with ParA to partition the ParB-*parS* nucleoprotein complex to each daughter cell. Here, we investigated the genome-wide distribution of ParB on the *Caulobacter* chromosome using a combination of *in vivo* chromatin immunoprecipitation (ChIP-seq) and *in vitro* DNA affinity purification with deep sequencing (IDAP-seq). We confirmed two previously identified *parS* sites and discovered at least three more sites that cluster ~8 kb from the origin of replication. We showed that *Caulobacter* ParB nucleates at *parS* sites and associates non-specifically with ~10 kb flanking DNA to form a high-order nucleoprotein complex on the left chromosomal arm. Lastly, using transposon mutagenesis coupled with deep sequencing (Tn-seq), we identified a ~500 kb region surrounding the native *parS* cluster that is tolerable to the insertion of a second *parS* cluster without severely affecting cell viability. Our results demonstrate that the genomic distribution of *parS* sites is highly restricted and is crucial for chromosome segregation in *Caulobacter*.

## INTRODUCTION

Proper chromosome segregation is essential in all living organisms if daughter cells are each to inherit a full copy of the genome. In eukaryotes, chromosome segregation during mitosis starts with sister chromosome condensation, followed by the formation of spindle fibres that attach to the kinetochore to pull sister chromatids apart. The kinetochore is the protein structure that assembles on the centromere and links each sister chromatid to microtubules polymers from the mitotic spindle. Unlike in eukaryotes, bacterial chromosome segregation happens without a dedicated spindle-like apparatus (1–3). Nevertheless, this process is highly organized and also involves protein-based components (4). The first segregated segment of the chromosome is usually proximal to the origin of replication (*ori*) (5–8). In many bacteria, this region is segregated by the tripartite ParA-ParB-*parS* partitioning system (6, 9–11). *parS* is a centromere-like DNA sequence that most often locates near *ori*. ParB is a DNA-binding protein that nucleates on a *parS* sequence. ParB is also capable of binding DNA non-specifically to spread along the chromosome from its cognate *parS* nucleation site (6, 12–14). Spreading was first discovered for the P1 plasmid-encoded ParB protein (15), and is subsequently found to be a general feature of many plasmid and chromosomal ParB proteins (13, 16–19). Spreading of AspA, a ParB-unrelated DNA segregation protein, has also been described for the archaeal *Sulfolobus* pNOB8 plasmid (20). ParB/Spo0J in a Gram-positive *Bacillus subtilis* might also bridge distal DNA together to coalesce into a large nucleoprotein complex (the “spreading and bridging” model) (12–14, 19). Similarly, the formation of the nucleoprotein complex for the F plasmid ParB-*parS* was proposed to happen via a “nucleation and caging” mechanism where the nucleation of ParB on *parS* creates a high local concentration of ParB, thereby caging ParB dimer-dimer together with non-specific DNA surrounding *parS* (21). Following ParB binding to *parS*, ParA, a Walker-box ATPase protein, interacts with ParB and powers the segregation of the ParB-DNA nucleoprotein complex to partition replicated chromosomes to each daughter cell (22, 23).

In *Caulobacter crescentus*, the ParA-ParB-*parS* system is essential for viability (11, 24). In G1-phase *Caulobacter*, *parS*/*ori* reside at one cell pole, the terminus (*ter*) is near the opposite pole, and the two chromosomal arms run orderly in parallel down the long axis of the cell (25, 26). After replication, the duplicated *parS* sites are released from the pole and separated slightly from one another before one *parS* site is translocated unidirectionally to the opposite cell pole. Toro *et al* (2008) identified two *parS* sites located ~8 kb from the *ori* on the left arm of the *Caulobacter* chromosome (8), while other works predicted six *parS* sites bioinformatically but did not report their sequences nor verify them experimentally (24, 27, 28). Furthermore, it is not yet known whether *Caulobacter* ParB spreads non-specifically on DNA, and if it does, how far it spreads along the chromosome from the *parS* nucleation site. Regarding the genome-wide distribution of *parS* sites, a comparative genomic study suggested that *parS* sites are not distributed randomly on bacterial chromosomes, rather they are found almost exclusively near the *ori* (7). Notably, in *Pseudomonas aeruginosa*, *parS* sites must be located within a ~650 kb region surrounding the *ori* for the chromosome segregation to proceed correctly (5).

In this study, we used genome-wide techniques (ChIP-seq and IDAP-seq) together with *in vitro* biochemical characterization to clarify the number and locations of *parS* sites in *Caulobacter*. We show that there are at least five *parS* sites clustered closely near the *ori* of *Caulobacter* chromosome, and that ParB occupies ~10 kb of DNA on the left arm of the chromosome. We also show that *Caulobacter* ParB nucleates on *parS* and spreads to flanking DNA independent of the location of *parS* on the chromosome. Moreover, using transposon mutagenesis coupled with deep sequencing (Tn-seq), we define a ~500 kb region surrounding the native *parS* cluster of the *Caulobacter* chromosome that is tolerable to the insertion of a second *parS* cluster without severely affecting cell viability. Our results demonstrate that the genomic location of *parS* is highly biased and crucial for proper chromosome segregation.

## MATERIALS AND METHODS

### Strains, media and growth conditions

*Escherichia coli* and *C. crescentus* were grown in LB and PYE, respectively. When appropriate, media were supplemented with antibiotics at the following concentrations (liquid/solid media for *C. crescentus*; liquid/solid media for *E. coli* [μg/mL]): carbenicilin (*E. coli* only: 50/100), chloramphenicol (1/2; 20/30), kanamycin (5/25; 30/50), spectinomycin (25/100; 50/50), oxytetracycline (1/2; 12/12), and apramycin (*E. coli* only: 25/50).

### Plasmids and strains construction

All strains used are listed in Supplementary Table S1. All plasmids and primers used in strain and plasmid construction are listed in Supplementary Table S2. For details on plasmids and strains construction, see the Supplementary Materials and Methods.

### Chromatin immunoprecipitation with deep sequencing (ChIP-seq)

*Caulobacter* cell cultures (25 mL) were grown in PYE and fixed with formaldehyde to a final concentration of 1%. Fixed cells were incubated at room temperature for 30 minutes, then quenched with 0.125 M glycine for 15 min at room temperature. Cells were washed three times with 1x PBS (pH 7.4) and resuspended in 1 mL of buffer 1 (20 mM K-HEPES pH 7.9, 50 mM KCl, 10% Glycerol and Roche EDTA-free protease inhibitors). Subsequently, the cell suspension was sonicated on ice using a probe-type sonicator (8 cycles, 15s ON, 15s OFF, at setting 8) to shear the chromatin to below 1 kb, and the cell debris was cleared by centrifugation (20 minutes at 13,000 rpm at 4°C).

The supernatant was then transferred to a new 2 mL tube and the buffer conditions were adjusted to 10 mM Tris-HCl pH 8, 150 mM NaCl and 0.1% NP-40. Fifty microliters of the supernatant were transferred to a separate tube for control (the INPUT fraction) and stored at −20ºC. In the meantime, antibodies-coupled beads were washed off storage buffers before adding to the above supernatant. We employed α-GFP antibodies coupled to sepharose beads (Abcam, UK) for ChIP-seq of CFP-ParB, α-FLAG antibodies coupled to agarose beads (Sigma, UK) for ChIP-seq of FLAG-ParB and FLAG-YFP, and Protein A beads (Sigma, UK) for α-ParB polyclonal antibody ChIP-seq of ParB. Briefly, 25 μL of beads was washed off storage buffer by repeated centrifugation and resuspension in IPP150 buffer (10 mM Tris-HCl pH 8, 150 mM NaCl and 0.1% NP-40). Beads were then introduced to the cleared supernatant and incubated with gentle shaking at 4°C overnight. In the next day, beads were then washed five times at 4°C for 2 min each with 1 mL of IPP150 buffer, then twice at 4°C for 2 min each in 1x TE buffer (10 mM Tris-HCl pH 8 and 1 mM EDTA). Protein-DNA complexes were then eluted twice from the beads by incubating the beads first with 150 μL of the elution buffer (50 mM Tris-HCl pH 8, 10 mM EDTA and 1% SDS) at 65°C for 15 min, then with 100 μL of 1X TE buffer + 1% SDS for another 15 min at 65°C. The supernatant (the ChIP fraction) was then separated from the beads and further incubated at 65°C overnight to completely reverse crosslink. The INPUT fraction was also de-crosslinked by incubation with 200 μL of 1X TE buffer + 1% SDS at 65°C overnight. DNA from the ChIP and INPUT fraction were then purified using the PCR purification kit (Qiagen) according to the manufacturer’s instruction, then eluted out in 50 μL of EB buffer (Qiagen). The purified DNA was then used directly for qPCR or being constructed into library suitable for Illumina sequencing using the NEXT Ultra library preparation kit (NEB). ChIP libraries were sequenced on the Illumina Hiseq 2500 at the Tufts University Genomics facility.

For *E. coli* ChIP-seq, cells harboring pUTC18-ParB (WT) or pUTC18-ParB (G101S) were grown in LB (50 mL) at 28°C to mid exponential phase (OD_600_ ~0.4) before 0.5 mM IPTG was added for an hour. Subsequently, formaldehyde is added to a final concentration of 1% to fix the cells. All following steps are identical to ChIP-seq for *Caulobacter*, except that we used α-T18 antibody coupled to sepharose beads (Abcam, UK) to immunoprecipitate ParB-DNA complexes.

For the list of ChIP-seq datasets in this study, see Supplementary Table S3.

### Generation and analysis of ChIP-seq profiles

For analysis of ChIP-seq data, Hiseq 2500 Illumina short reads (50 bp) were mapped back to the *Caulobacter* NA1000 reference genome (NCBI Reference Sequence: NC*-*011916.1) using Bowtie 1 (29) and the following command:

~~~
bowtie ‐m 1 ‐n 1 ‐‐best ‐‐strata ‐p 4 ‐‐chunkmbs 512 NA1000*-*bowtie ‐‐sam *.fastq > output.sam
~~~

Subsequently, the sequencing coverage at each nucleotide position was computed using BEDTools (30) using the following command:

~~~
bedtools genomecov ‐d ‐ibam output.sorted.bam ‐g NA1000.fna > coverage_output.txt
~~~

For analysis of *E. coli* ChIP-seq data, reference genomes were first reconstructed *in silico* by inserting the nucleotide sequence of *parS* and apramycin antibiotic resistance cassette to the *ybbD* locus of *E. coli* MG1655 genome. Afterwards, Hiseq 2500 Illumina short reads were mapped back to these reconstructed reference genomes using Bowtie 1. Sequence coverage at each nucleotide position was also computed using BEDTools. Finally, ChIP-seq profiles were plotted with the x-axis representing genomic positions and the y-axis is the number of reads per base pair per million mapped reads (RPBPM) or number of reads per kb per million mapped reads (RPKPM) using custom R scripts.

### *In vitro* DNA affinity purification with deep sequencing (IDAP-seq)

*Caulobacter* genomic DNA was fragmented using a Diagenode Bioruptor to 200 bp-500 bp in length. Five μg of genomic DNA was incubated with 320 nM of purified ParB-(His)_6_ in IDAP buffer (20 mM K-HEPES pH7.9, 50 mM KCl, 10 % Glycerol, 10 mM Tris pH 8, 150 mM NaCl, 0.1% (v/v) Surfactant P20) at room temperature. After 60 minute incubation at room temperature, 100 μL of Cu^2+^ Talon Superflow beads (GE Healthcare) were added, and the mixture was left at 4°C with gentle shaking for a further 60 minutes. Afterwards, Talon beads were repeatedly washed in IPP150 buffer (10 mM Tris pH 8, 150 mM NaCl, 0.1% NP40) and 1xTE buffer (10mM Tris pH 7.4, 1 mM EDTA) to wash off unbound ParB. ParB-DNA complexes were then eluted from the beads by incubating the beads with 150 μL of the elution buffer (50 mM Tris-HCl pH 8, 10 mM EDTA and 1% SDS) at 65°C for 15 min, then with 100 μL of 1X TE buffer + 1% SDS for another 15 min at 65°C. Subsequently, DNA was purified using a Qiaquick PCR clean up kit before being made into a library suitable for Illumina sequencing using the NEXT Ultra library preparation kit (NEB). IDAP-seq libraries were sequenced on the Illumina Hiseq 2500 at the Tufts University Genomics facility. As a control, Talon beads were also incubated with fragmented genomic DNA in the absence of ParB-(His)_6_. Eluted DNA from the negative control was also made into Illumina sequencing library and sequenced in parallel to control for DNA fragments that bind to the surface of Talon beads non-specifically.

### Analysis of IDAP-seq data to pinpoint *parS* sites to a single-nucleotide resolution

For analysis of IDAP-seq data, Hiseq 2500 Illumina short reads (50 bp) were mapped back to the *Caulobacter* NA1000 reference genome (NCBI Reference Sequence: NC*-*011916.1) using Bowtie 1 (29) and the following command:

~~~
bowtie ‐m 1 ‐n 1 ‐‐best ‐‐strata ‐p 4 ‐‐chunkmbs 512 NA1000*-*bowtie ‐‐sam *.fastq > output.sam
~~~

Subsequently, sequencing reads were sorted to either being mapped to the upper DNA strand or to the lower strand of the reference genome, as suggested in the original IDAP-seq publication (31). The number of 5’ end of reads that were mapped to the upper strand was counted for each nucleotide position along the *Caulobacter* genome using BEDTools (30) and the following command:

~~~
bedtools genomecov ‐d ‐5 ‐strand + ‐ibam output.sorted.bam ‐g NA1000.fna > upper_strand_output.txt
~~~

To count the number of 5’ end of reads that were mapped to the lower strand, the following command was used instead:

~~~
bedtools genomecov ‐d ‐5 ‐strand - ‐ibam output.sorted.bam ‐g NA1000.fna > lower_strand_output.txt
~~~

The IDAP-seq profile was then plotted using R. The sequence in between the summit of upper strand profile and that of the lower strand profile defines the minimal *parS* sequence required for binding to ParB. See also Fig. S3 for the principle behind the strand-specific analysis of IDAP-seq data to determine DNA-binding sequence at nucleotide resolution.

### Transposon mutagenesis coupled with next-generation sequencing (Tn-seq)

The Tn5 transposon delivery plasmid (pMCS1-Tn5-ME-R6Kγ-kan^R^-ME or pMCS1-Tn5-ME-R6Kμ-kan^R^-*parS*^345^-ME) was conjugated from an *E. coli* S17-1 donor into *Caulobacter* cells. Briefly, *E. coli* S17-1 was transformed with the transposon delivery plasmid and plated out on LB + kanamycin. On the next day, colonies forming on LB + kanamycin were scraped off the plates and resuspended in PYE to OD_600_ of 1.0. Cells were pelleted down and resuspended in fresh PYE twice to wash off residual antibiotics. 100 μL of cells were mixed with 1000 μL of exponentially growing *Caulobacter* (either wild-type, *Δsmc*, Flip 1-5, or Flip 2-5 *Caulobacter* cells), then the mixture was centrifuged at 13,000 rpm for 1 minute. The cell pellet was subsequently resuspended in 50 μL of fresh PYE and spotted on a nitrocellulose membrane resting on a fresh PYE plates. Twenty conjugations were performed to generate Tn5 insertion library for each *Caulobacter* strain. PYE plates with nitrocellulose disks were incubated at 30°C for 5 hours before being resuspended by vortexing vigorously in fresh PYE liquid to release bacteria. Resuspended cells were plated out on twenty 30 cmx30 cm square Petri disks containing PYE agar supplemented with kanamycin and carbenicilin, and incubated for 3 days at 30°C. After 3-day incubation, cells (~500,000-1,000,000 single colonies) were scraped off the Petri disk and resuspended in 200 mL of fresh PYE. The culture was pipetted repeatedly using a 10 mL glass pipette to break clumps and homogenize the culture. Genomic DNA was subsequently extracted from a 2 mL sample using a genomic DNA extraction kit (Qiagen). Genomic DNA (1 μg) was sheared to between 200 bp and 500 bp using a Diagenode Bioruptor Plus (30 sec ON, 30 sec OFF, for 20 cycles at low sonication power). The fragmented DNA were resolved on a 2% agarose gel and a band of desired DNA length (200 bp-500 bp) was excised and extracted using a QiaQuick gel extraction kit (Qiagen) before being made into an Illumina deep sequencing libraries.

For the list of Tn-seq libraries in this study, see Supplementary Table S3. For details on the construction of Illumina libraries, see Supplementary Materials and Methods.

### Analysis of Tn-seq data

Hiseq 2500 Illumina short reads (50 bp) were mapped back to the *Caulobacter* NA1000 reference genome (NCBI Reference Sequence: NC*-*011916.1) using Bowtie 1 (29) and the following command:

~~~
bowtie ‐m 1 ‐n 1 ‐‐best ‐‐strata ‐p 4 ‐‐chunkmbs 512 NA1000*-*bowtie ‐‐sam *.fastq > output.sam
~~~

For *Caulobacter* strains with an inverted DNA segment, a reconstructed fasta file with the correct orientation for the inverted segment was used as reference genome for Bowtie instead. Subsequently, the sequencing coverage for each nucleotide position was computed using BEDTools (30) and the following command:

~~~
bedtools genomecov ‐d ‐ibam output.sorted.bam ‐g NA1000.fna > coverage_output.txt
~~~

Finally, the ratio between the number of reads of libraries generated from pMCS1-Tn5-ME-R6Kγ-kan^R^-ME or pMCS1-Tn5-ME-R6Kγ-kan^R^-*parS^4+5+6^*-ME were calculated. Results were binned over 10 kb and represented as a log_10_ scale.

### Measure ParB-*parS* binding affinity by Surface Plasmon Resonance (SPR)

Single-stranded oligomers containing *parS* sequence were purchased from Sigma and reconstituted to 100 μM in water. Complementary oligos were annealed together in an annealing buffer (10 mM Tris-HCl pH 8.0, 50 mM NaCl, and 1 mM EDTA) to form double stranded DNA before being diluted to a working concentration of 1 μM in HPS-EP+ buffer (0.01 M HEPES pH 7.4, 0.15 M NaCl, 3 mM EDTA, 0.005% v/v Surfactant P20) for each SPR experiment. The sequences of DNA oligos used in this study are reported in Supplementary Table S2. SPR measurements were recorded at 25°C using a Biacore T200 system (GE Healthcare). All experiments were performed using *R*e-usable *D*NA *Ca*pture *T*echnique (ReDCaT) exactly as described in (32). Briefly, ReDCAT uses a Sensor Chip SA (GE Healthcare), which has streptavidin pre-immobilized to a carboxymethylated dextran matrix, to which a 20 base biotinylated ReDCaT linker is immobilised. This is then used to immobilize *parS*-containing biotin-labelled double stranded oligos on the chip surface as each contain a single stranded overhand complimentary to the ReDCaT linker on the surface. The DNA to be tested is flowed over one flow cell on the chip at a flow rate of 10 μl/ min and it anneals through the complementary DNA to the ReDCaT linker. *C. crescentus* ParB-(His)_6_ or *B. subtilis* Spo0J-(His)_6_, pre-diluted in HBS-EP+ buffer, was then flowed over the chip surface (the blank surface and the one with the DNA immobilised) and then HBS-EP+ buffer was then passed over to allow ParB-(His)_6_ to dissociate from DNA. A high-salt wash buffer was injected to the chip to wash off any residual ParB-(His)_6_ protein on the chip’s surface. The test DNA could then be removed using a wash with 1M NaCl, 50mM NaOH. The chip could then be used again to load a new piece of test DNA. The SPR signal (Response Units) was monitored continuously throughout the process. Each cycle was repeated for increasing concentrations of ParB-(His)_6_. For each concentration, the amount of ParB bound was measured and plotted against the concentration to construct a ParB-parS binding curve (Fig. S2). All sensorgrams recorded during ReDCAT experiments were analyzed using Biacore T200 BiaEvaluation software version 1.0 (GE Healthcare). Data were then plotted using Microsoft Excel or R, and K_d_ was estimated from best-fit curves.

### Fluorescence microscopy image analysis

*C. crescentus* strain MT190 or strains with ectopic *parS^3+4^* (at +200 kb, +1000 kb or +1800 kb) were grown to OD_600_=0.4 in the presence of appropriate antibiotics before being spotted to agarose pad for microscopy observation. Phase contrast (150 ms exposure) and fluorescence images (1000 ms exposure) were collected. MicrobeTracker (http://microtracker.org) was used to detect cell outlines and cell length (33). SpotFinderM was used to manually detect fluorescent foci positions (33). Data (cell length, foci number) were exported to ․csv files and subsequently analyzed and plotted in R.

## RESULTS AND DISCUSSION

### ParB occupies a 10 kb DNA region near the origin of replication

To define the distribution of ParB on the chromosome, we performed chromatin immunoprecipitation with deep sequencing. We fused the *flag* tag to the ParB-encoding gene at its 5’ end and placed this allele downstream of a vanillate-inducible promoter (P*_van_*), at the chromosomal *vanA* locus. The vanillate-inducible *flag-parB* was then transduced to a *Caulobacter* strain where the native and untagged *parB* was under the control of a xylose-inducible promoter (P*_xyl_*). *Caulobacter* cells were depleted of untagged ParB by addition of glucose for 5 hours, then vanillate was added for an additional hour before cells were fixed with 1% formadehyde for ChIP-seq (Fig. 1A). *Caulobacter* cells depleted of native ParB while producing the FLAG-tagged ParB version are viable, indicating that the tag does not interfere with ParB function (Fig. S1A). For ChIP-seq, DNA-bound to FLAG-ParB was pulled down using α-FLAG antibody coupled to sepharose beads. The immunoprecipitated DNA was deep sequenced and mapped back to the *Caulobacter* genome to reveal enriched genomic sites (Fig. 1A). As a negative control, we performed α-FLAG ChIP-seq in a *Caulobacter* strain that produces FLAG-tagged YFP, a non-DNA binding protein (Fig. 1B). The ChIP-seq profile of FLAG-ParB showed a clear enrichment in the DNA region on the left chromosomal arm, ~8 kb away from the origin of replication. No other significant enrichment was observed elsewhere on the chromosome or in the negative control (Fig. 1A-B). A closer examination of the *ori*-proximal region revealed an extended ~10 kb region with significant enrichment above background and four defined peaks (Fig. 1A). To independently verify our results, we repeated the ChIP-seq experiment using α-GFP antibody to pull down DNA from a *Caulobacter* strain that produces a CFP-ParB fusion protein from its native location as the only source of ParB in the cell or using a polyclonal α-ParB in a wild-type *Caulobacter* (Fig. S1B). For all cases, we retrieved very similar ChIP-seq profiles to that of FLAG-ParB, suggesting the extended DNA region associating with ParB is not an artefact of tagging but a property of *Caulobacter* ParB itself.

**Figure 1.**
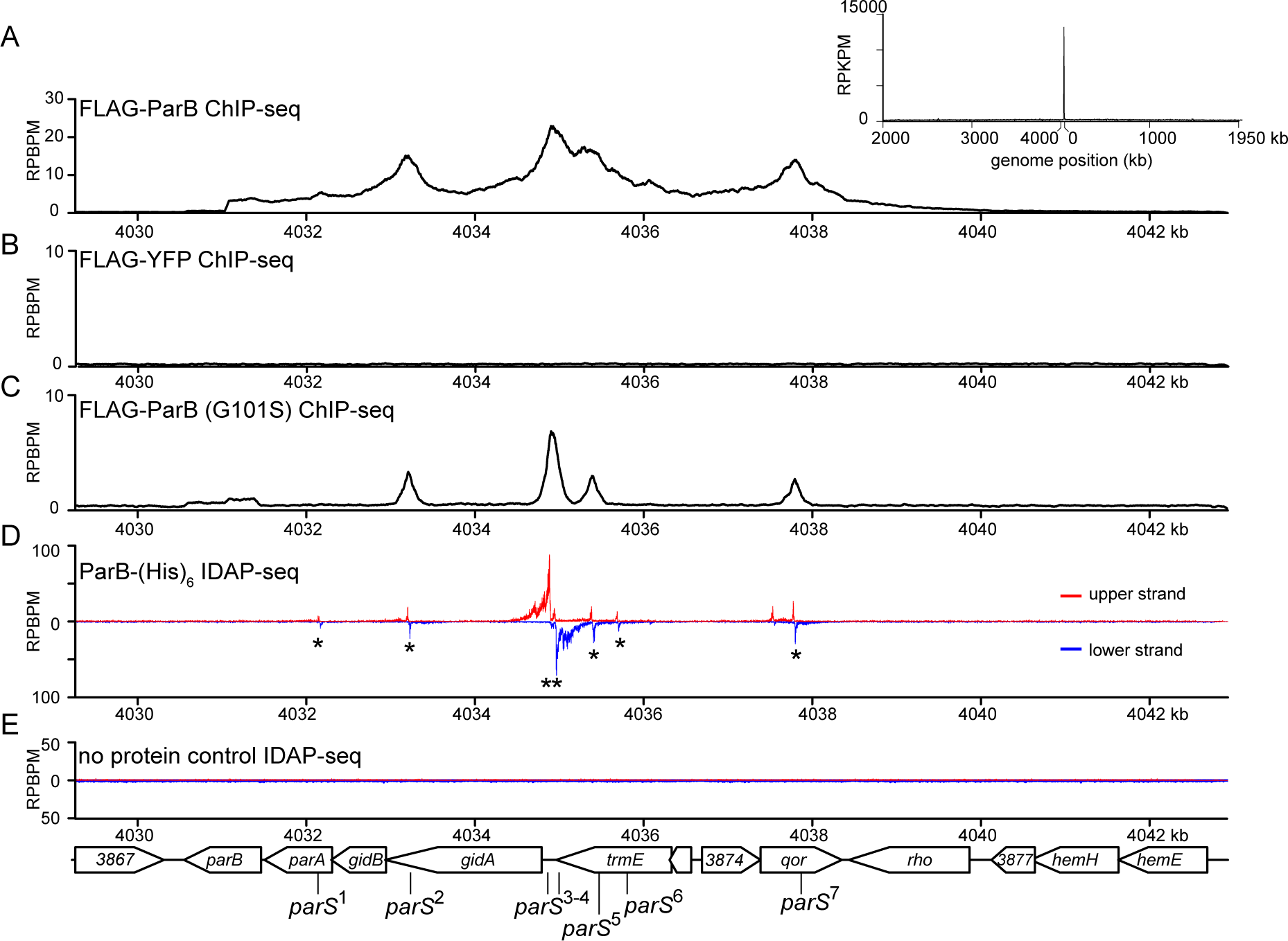
ParB occupies 10 kb DNA region near the origin of replication. **(A)** The distribution of FLAG-tagged ParB on *Caulobacter* chromosome between +4030 kb and +4042 kb. ChIP-seq signals were reported as the number of reads at every nucleotide along the genome (RPBPM value). The whole-genome ChIP-seq profile of ParB is shown in the inset. For the whole genome profile, the ChIP-seq signals were reported as the number of reads at every kb along the genome (RPKPM value). **(B)** ChIP-seq profile of FLAG-tagged YFP. **(C)** ChIP-seq profile of FLAG-tagged ParB (G101S) mutant. **(D)** IDAP-seq profile of ParB-(His)6 with sonication-fragmented genomic DNA from *Caulobacter*. IDAP-seq reads were sorted to either the upper strand (red) or to the lower strand (blue) of the reference genome to enable identification of *parS* sites (see also Fig. 2 and Fig. S3). Putative *parS* sites (1 to 7) are noted with asterisks (see also Fig. 2). **(E)** IDAP-seq profile of a negative control in which ParB-(His)_6_ was omitted.

The extensive 10-kb ParB-binding DNA region cannot be explained by the length of DNA fragments that were sheared as part of a ChIP-seq protocol. We sequenced immunoprecipitated DNA from both ends to determine their exact size distribution (Table S3). Pulled-down DNA averages around 150 bp, much smaller than the size of ChIP-seq peaks in our study. However, the extended ParB-binding DNA region can be most easily explained by the non-specific binding of ParB to DNA outside of the *parS* nucleation site, either by a “spreading and bridging” or “caging” mechanism. If so, *Caulobacter* ParB mutants that are impaired in binding to non-specific DNA are predicted to spread less. To identify such mutants in *Caulobacter*, we mutated the highly-conserved N-terminal Box II motif which was shown to be important for the non-specific DNA-binding activity of *B. subtilis* ParB (Fig. S2A) (12, 19). Four variants were constructed *parB* (G101S), *parB* (R103A), *parB* (R104A), and *parB* (R106A). We introduced the *flag*-tagged *parB* mutant allele at the *van* locus, in the P*_xyl_-parB* genetic background, then employed α-FLAG ChIP-seq to assess the distribution of mutated ParB on the chromosome. Two mutants, ParB (G101S) and ParB (R104A), were found to produce well-defined and symmetrical peaks (~400 bp in width) that are typical of site-specific DNA-binding proteins (Fig. 1C and Fig. S1B). On the contrary, wild-type ParB peaks are much wider and asymmetrical (Fig. 1A). These data suggest that *Caulobacter* ParB, similar to *B. subtilis* and *P. aeruginosa* ParB, also spreads along the chromosome. Lastly, we noted that DNA enrichment in ChIP-seq experiments with ParB (G101S) or ParB (R104A) is ~5 fold less than that of wild-type ParB (Fig. 1A-C), despite the fact that ParB variants nucleate equally well on DNA *in vitro* (Fig. S2B). This is most likely because ParB (G101S) and ParB (R104A) are less stable than wild-type ParB *in vivo* (Fig. S2C).

### Identification of *parS* sites and correlating ParB-*parS in vitro* binding affinities to their *in vivo* ChIP-seq enrichment

Since the large width of ChIP-seq peaks obscures the exact position of *parS*, we employed *in vitro* DNA affinity purification with deep sequencing (IDAP-seq) (31) to pinpoint *parS* sequence to near single-nucleotide resolution. Purified ParB-(His)6 was incubated with randomly-fragmented *Caulobacter* genomic DNA, then ParB-DNA complexes were pulled-down using immobilized Ni^2+^ beads. ParB-bound DNA fragments were eluted out and sequenced *en masse*. The sequencing reads were mapped back to either the upper strand or the lower strand of the *Caulobacter* genome (Fig. 1 D and Fig. 2). Analysis of the strand-specific coverage map allows identification of seven 16 bp putative *parS* sites (see Fig. 1D and Fig. S3 for the methodology of IDAP-seq data analysis). These included the two *parS* sites (sites 3 and site 4) that were first discovered in Toro *et al* (2008) (8) but revealed five more putative sites (sites 1, 2, 5, 6 and 7).

**Figure 2.**
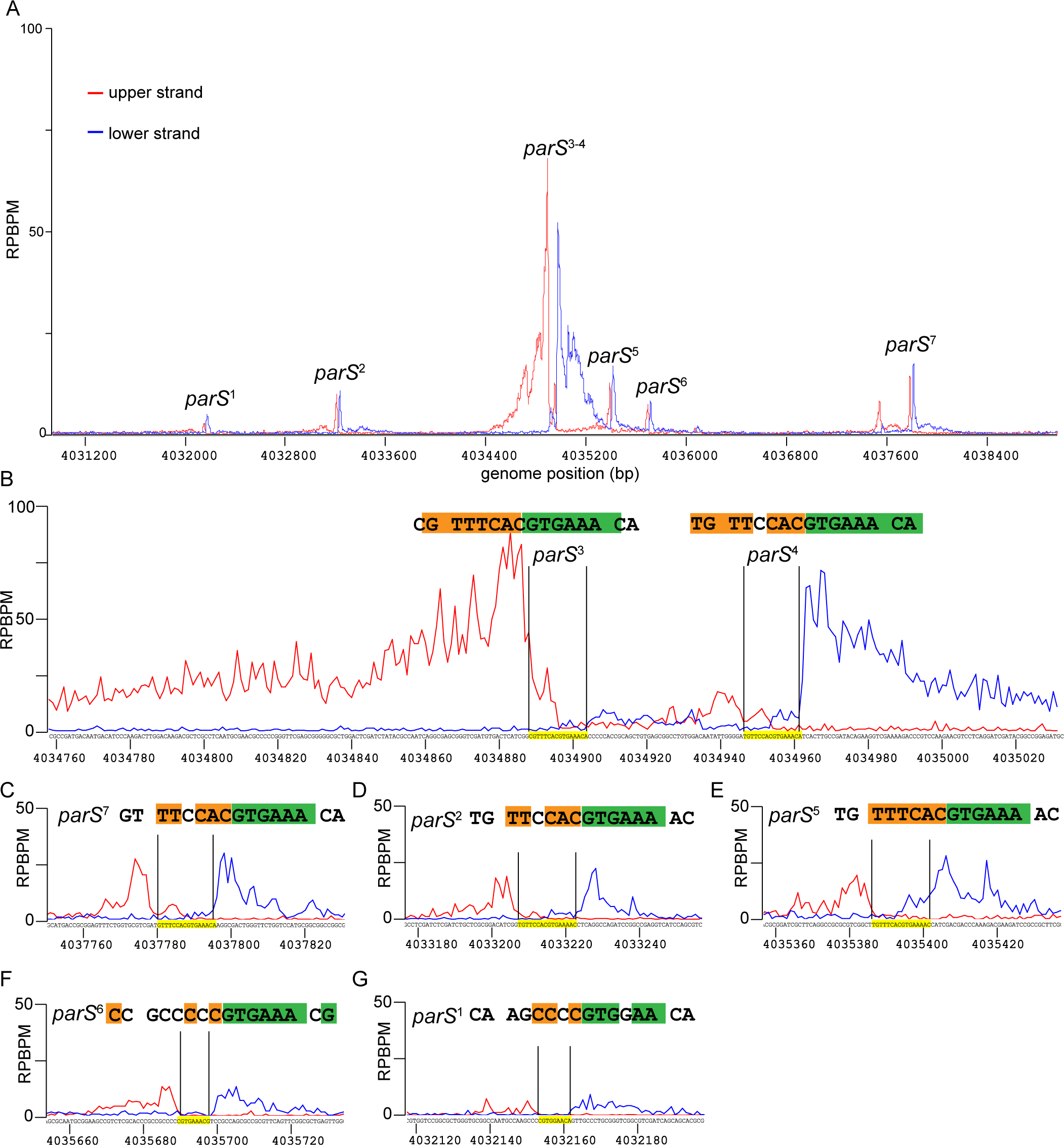
Identification of *parS* sequences by *in vitro* DNA purification with deep sequencing (IDAP-seq) Sequencing reads were sorted to either the upper DNA strand (red) or to the lower strand (blue) of the *Caulobacter* reference gnome, as suggested in the original IDAP-seq publication (Belitsky and Sonenshein, 2013, Fig. S3). The sequence in between the summit of the upper strand profile and that of the lower strand profile defines the *parS* sequence required for binding to ParB *in vitro* (See also Fig. S3). **(A)** IDAP-seq profile of ParB-(His)_6_ in the genomic region between +4031 kb and +4039 kb. **(B-G)** IDAP-seq profile of ParB-(His)_6_ surrounding each individual *parS* site. Palindromic nucleotides within the identified *parS* site are shaded in orange and green.

To correlate the sequence conservation to the binding affinity of ParB, we measured the equilibrium dissociation constant (K_d_) of ParB binding to 24-bp double-stranded oligonucleotides containing individual putative *parS* sites by Surface Plasmon Resonance (SPR) (Fig. 3 and Fig. S4). The double-stranded oligonucleotides was tethered to a chip surface within an SPR flow cell. Purified ParB-(His)_6_ was flowed over the test DNA. ParB binding was recorded by measuring the change in response units during ParB injection. After injection, the chip was washed with buffer and subsequently with high salt buffer to remove any bound ParB. This cycle was repeated for an increasing concentration of ParB dimer to enable the estimation of K_d_ (Fig. 3 and Fig. S4). Note that the length of the double-stranded oligonucleotides was limited to 24 bp so that only the nucleating event of ParB on *parS* was observed, and not the interaction with DNA flanking *parS*. We observed that sites 2, 3, 4, 5, and 7 have low nM K_d_ values (Fig. 3), consistent with their high ChIP-seq peaks (Fig. 1). On the other hand, ParB binds to the putative sites 1 and 6 weakly *in vitro*, albeit more than to a scrambled *parS* control (Fig. 3), suggesting that sites 1 and 6 are perhaps unlikely to be significant *in vivo*.

**Figure 3.**
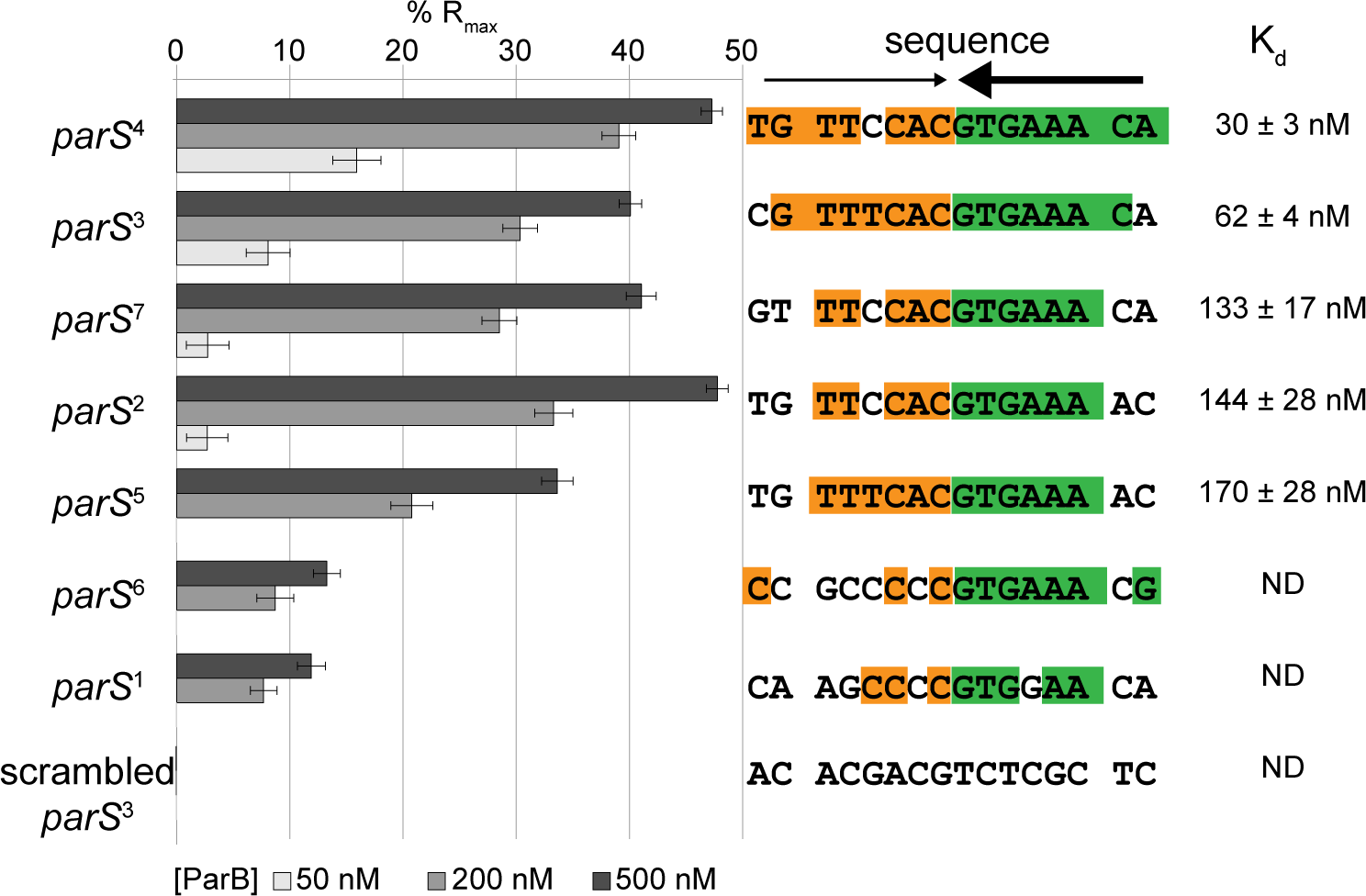
ParB-*parS in vitro* binding affinities correlate to their *in vivo* ChIP-seq enrichment. Surface Plasmon Resonance (SPR) was used to measure binding affinity of ParB (50 nM, 200 nM and 500 nM) to 24-bp double-stranded DNA that contains individual putative *parS* site. The level of ParB binding to DNA was expressed as a percentage of the theoretical maximum response, R_max_, assuming a single ParB dimer binding to one immobilized double-stranded DNA oligomer. This normalization process enabled the various responses to be readily compared, irrespective of the quantity length of the DNA tethered on an SPR chip surface. A wider range of ParB concentration (6.25 nM, 12.5 nM, 25 nM, 50 nM, 100 nM, 200 nM, 400 nM, 600 nM and 800 nM) was used to estimate the binding constant (K_d_) of ParB to individual *parS* site (Fig. S4). The sequences of *parS* are shown with palindromic nucleotides shaded in orange and green. Convergent arrows on top of *parS* sequence indicate that *parS* sites are palindromic. Thicker arrow signifies that the second half of *parS* sequences (GTGAAA, in green) is conserved among *Caulobacter parS* sites.

Importantly, the affinity of *Caulobacter* ParB for its *parS* site (30 nM ± 3 nM) is much stronger than the previously reported K_d_ for the *B. subtilis* Spo0J-*parS* interaction (230 ± 7 nM) (6, 14). To check whether the difference in K_d_ is due to measurement techniques, we purified *B. subtillis* Spo0J and determined its affinity to a cognate *parS* or to a randomized site by SPR (Fig. S5). We found that the apparent K_d_ for *B. subtilis* Spo0J-cognate *parS* is 114 ± 21 nM, and *B. subtilis* Spo0J-randomized *parS* is 183 ± 29 nM (Fig. S5). These values are in a similar range to those measured previously using a different technique (14). Our experiments also confirmed the previous finding that *B. subtilis* Spo0J does not discriminate well between *parS* and non-*parS* DNA (14). Based on the similar Kd for *parS* and non-*parS* site, it has been suggested that the presence of *parS* site does not promote non-specific DNA binding and/or condensation events by *B. subtilis* Spo0J (14). On the contrary, *Caulobacter* ParB binds *parS* tightly but almost does not bind or binds very weakly to non-*parS* site (Fig. 3 and Fig. S4). Nevertheless, *in vivo* ChIP-seq experiments showed unequivocally that *Caulobacter* ParB spreads to non-specific DNA on both sides of the core *parS* sequence (Fig. 1 and Fig. S1). Our results with *Caulobacter* ParB, therefore, support the idea that the initial ParB-*parS* nucleation event is important for spreading. Why is there a stark contrast between the two ParB proteins of the same class? Recently, it has been showed that the C-terminal domain of *B. subtilis* Spo0J, in addition to the middle helix-turn-helix domain, binds DNA non-specifically and contributes to DNA condensation (14 and M. Dillingham, personal communications). In *Caulobacter*, the C-terminal domain of ParB is not similar to that of the *B. subtilis* Spo0J, hence might not bind non-specific DNA strongly. The DNA-binding property of the Spo0J C-terminal domain might explain why *Bacillus parS* sites do not cluster as closely as in *Caulobacter*. The four strongest *Bacillus parS* sites (*parS* at 354°, *parS* at 355°, *parS* at 356°, and *parS* at 359°) are ~5 kb, 13 kb, and 39 kb apart from each other, respectively. On the contrary, the five strongest *Caulobacter parS* sites are all within a 5-kb DNA segment. The lower capability of *Caulobacter* ParB in binding to non-specific DNA might necessitate a closer clustering of *parS* sites for an efficient “spreading” in this bacterium. We explore this possibility by investigating the spreading of *Caulobacter* ParB from individual *parS* sites below.

### ParB spreads to a maximum of 2 kb around individual *parS* site

Since *parS* sites are located within essential genes or genes that have a high fitness cost, we were not able to ablate individual *parS* sites to investigate the spreading of ParB in *Caulobacter*. Instead, we investigated the spreading of ParB from individual *parS* sites by expressing the *Caulobacter* ParB/*parS* system in *E. coli*. Since *E. coli* does not possess a ParB homolog nor a *Caulobacter parS*-like sequence, it serves as a suitable heterologous host for this experiment. We inserted individual *parS* sites onto the *E. coli* chromosome at the *ybbD* locus (Fig. 4). The ParB protein was expressed from an IPTG-inducible promoter as a C-terminal fusion to the T18 fragment of *Bordetella pertussis* adenylate cyclase. The T18-ParB is fully functional in *E. coli* as judged by its interactions with their known partners such as ParB itself, ParA, and MipZ in a bacterial-two hybrid assay (Fig. S6A). We induced exponentially-growing *E. coli* cells at 28°C with 500 μM IPTG for an hour before fixing with formadehyde for ChIP-seq. DNA bound to T18-ParB was immunoprecipitated using α-T18 conjugated sepharose beads. A scrambled *parS* site 3 was also inserted at the *ybbD* locus to serve as a negative control. As expected, the strong *parS* sites (sites 2, 3, 4, 5, and 7), on their own showed a high level of DNA enrichment, in agreement with their *in vitro* ParB binding affinity (Fig. 4). The weak putative *parS* sites (site 1 and 6) show little to no enrichment above background (Fig. S6B). Most importantly, we observed that ParB in an *E. coli* host spreads to a maximum of ~2 kb around each *parS* site (Fig. 4), much less than ~10 kb for *B. subtilis* Spo0J-single *parS* (12). Next, we repeated the ChIP-seq experiment but with a spreading-defective ParB (G101S). This revealed symmetrical peaks with a ~400-bp width, confirming that *Caulobacter* ParB can spread to any neighbouring DNA and that non-specific interaction with DNA is mainly dependent on an initial ParB-*parS* nucleation event. Lastly, we noted that the spreading of wild-type ParB is not equal on both sides of *parS*. It is likely that the non-specific association of ParB with neighbouring DNA might be influenced by on-going transcription or other nearby DNA-binding proteins. This asymmetrical spreading has been observed previously with ParB homologs from other bacterial species (19, 34).

**Figure 4.**
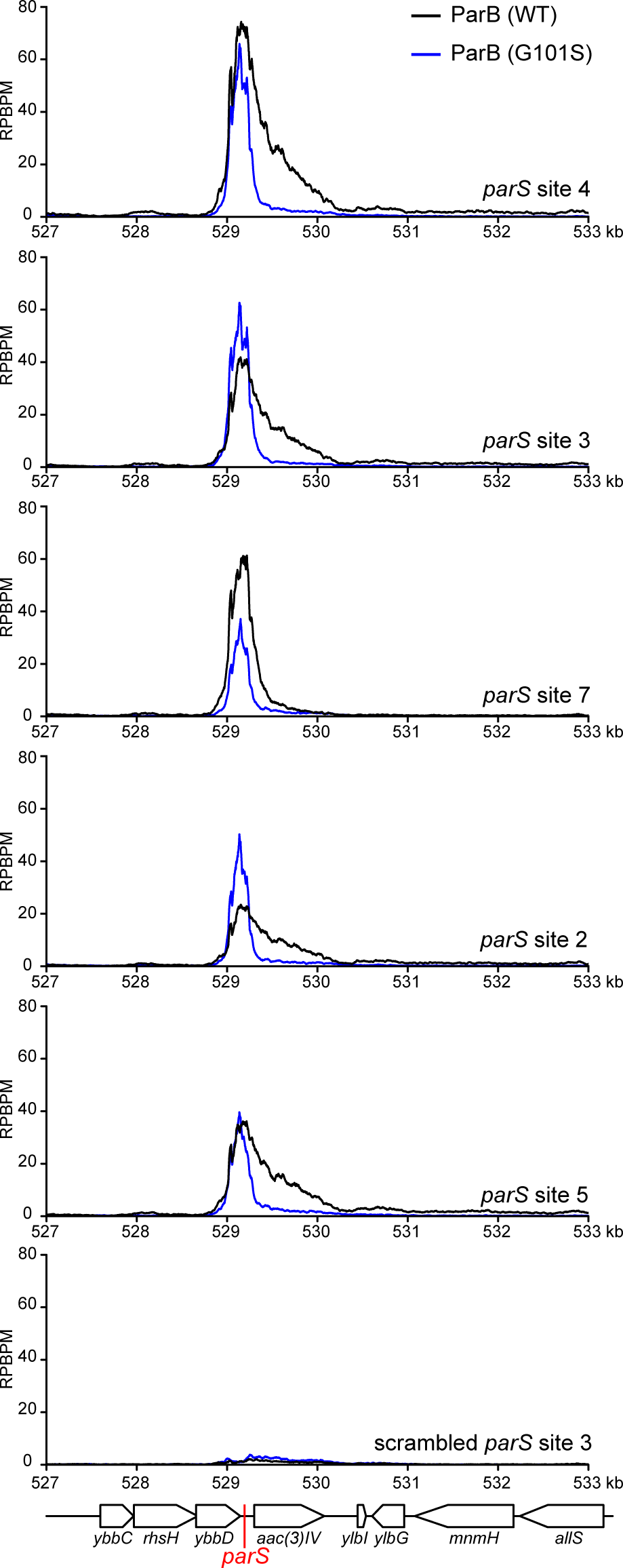
*Caulobacter* ParB binds to *parS* and spreads to flanking DNA in a heterologous *E. coli* host. A cassette composed of individual *parS* (red line) site and an apramycin resistance marker *aac(3)IV* was inserted at the *yybD* locus on an *E. coli* chromosome. T18-ParB (WT) (black) or T18-ParB (G101S) (blue) were expressed from an IPTG-inducible promoter, and their distribution on the *E. coli* chromosome were determined by α-T18 ChIP-seq. ChIP-seq signals were reported as the number of reads at every nucleotide along the genome (RPBPM value). A cassette composed of a scrambled *parS* site 3 and an apramycin resistance marker was also inserted at the *yybD* locus and serves as a negative control.

Since *Caulobacter* ParB associates maximally with only ~2 kb DNA surrounding individual *parS* site, the clustering of *parS* sites might serve to enable a higher concentration of DNA-bound ParB near *ori* than is possible with a single site. A previously study estimated that ~80% of the total cellular ParB is bound at *parS* sites in *Caulobacter* (1). *Caulobacter* ParA was also found to require a higher concentration of DNA-bound ParB than in *B. subtilis* to activate its ATPase activity, an essential step for chromosome segregation by the ParAB-*parS* system (1). Furthermore, it is known that *Caulobacter* ParB interacts with MipZ, which in turns binds PopZ to anchor the *ori*-proximal DNA to the cell pole (35–37). A high local concentration of DNA-bound ParB would enable a robust anchorage of the *ori* DNA domain to the cell pole. We noted that the nucleation-competent but spreading-defective ParB (G101S) or ParB (R104A) variants are unable to support *Caulobacter* growth, implying that ParB spreading is required for cell viability (Fig. S1A). In line with our study, *B. subtilis* or *P. aeruginosa* engineered with a single *parS* are defective in chromosome segregation, resulting in elevated numbers of anucleate cells (5, 19, 38).

### Extra copies of *parS* can reduce the fitness of *Caulobacter* depending on their genomic locations

Additional copies of *parS*, for example when is placed on a multi-copy number plasmid, can be lethal for cells because plasmid DNA can be segregated instead of the chromosome, resulting in daughter cells with either zero or two chromosomes (8). Indeed, we found the presence of a *parS*-carrying plasmid caused growth impairment in *Caulobacter*, and the fitness cost correlates well with the ParB-*parS* binding affinity (Fig. 5). Plasmid-borne sites 3 and 4, which are the strongest *parS* sites, reduced cell viability by ~1000 fold compared to a negative control (scrambled site 3). Extra copies of sites 2, 5 and 7 reduced cell viability by ~100 fold compared to a control, while the weaker *parS* sites 1 and 6 did not impact cell viability when present on a plasmid.

**Figure 5.**
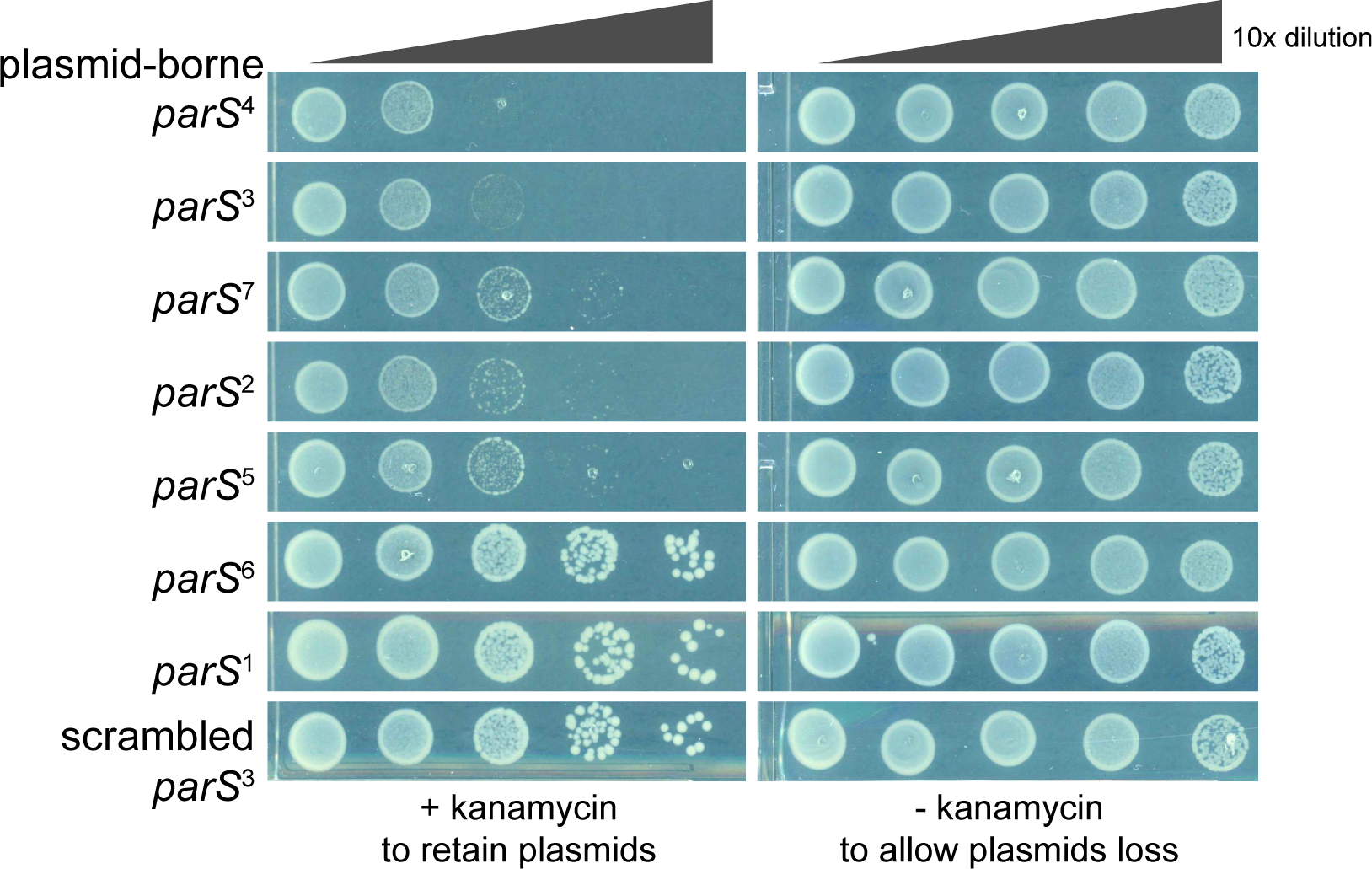
Plasmid-borne *parS* reduces the fitness of *Caulobacter*. Low-copy number plasmid harbouring individual *parS* site was conjugated from *E. coli* S17-1 to wild-type *Caulobacter.* The same number of *E. coli* and *Caulobacter* cells were used for each conjugation. A ten-fold serial dilution was performed and spotted on PYE plates supplemented with both nalidixic acid and kanamycin or just with nalidixic acid. Addition of kanamycin enforces the retention of *parS* plasmid, while omitting kanamycin allows plasmid loss. All cells were spotted on the same +kanamycin or ‐kanamycin plates, and pictures were taken after 3 day incubation at 30°C.

We reasoned that if the toxicity of a plasmid-borne *parS* site was due to the segregation of plasmids instead of the chromosome then having extra *parS* sites on the chromosome should eliminate the toxicity. Indeed, we were able to engineer a 260-bp DNA segment containing both strong *parS* site 3 and site 4 at various positions from *ori* to *ter* on both arms of *Caulobacter* chromosome. On the contrary, a plasmid containing both *parS* sites 3 and 4 is completely lethal to *Caulobacter* cells (8). Nevertheless, we noted a variation in the fitness of *Caulobacter* with extra chromosomal *parS* sites, depending on the location of the ectopic *parS* (Fig. 6). An extra *parS*^3+4^ inserted at +200 kb (near *ori*) or at +1800 kb (near *ter*) did not impact the fitness of the cell dramatically as judged by a normal cell length distribution and a 6-fold increase in the number of anucleate cells (Fig. 6B and Fig. 6D). On the contrary, *parS*^3+4^ inserted at +1000 kb (middle of the right arm of the chromosome) caused a more severe fitness defect. The cells were more elongated (4.74 ± 3.3 μm) compared to WT (2.97 ± 0.77 μm) (Fig. 6). Furthermore, the number of cells with no or more than two CFP-ParB foci were elevated ~ 11 fold in comparison to strains without an ectopic *parS*^3+4^ (Fig. 6C). Lastly, in *Caulobacter*, ParB recruits MipZ, which in turns regulates the positioning of the division plane (37). We found that the number of MipZ-CFP foci are abnormal in strains with an ectopic *parS*^3+4^ site, suggesting that cell division defects also contribute to a lower cell fitness in those strains (Fig. S7). Taken all together, our data suggest that the genomic location of an extra chromosomal copy of *parS* is important for the cell fitness in *Caulobacter*.

**Figure 6.**
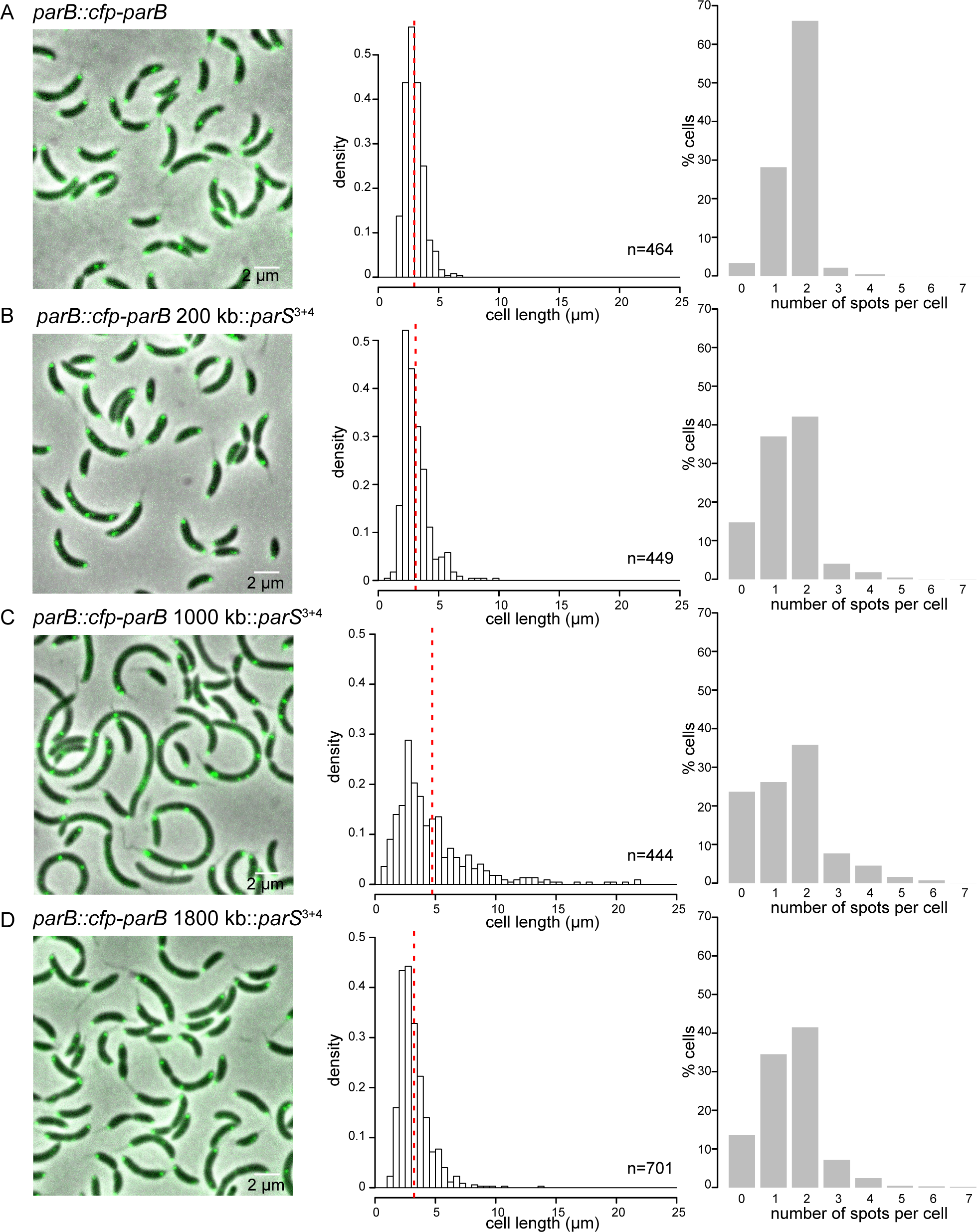
The position of an ectopic *parS* on the chromosome is critical for the fitness of *Caulobacter*. Micrograph of *parB::cfp-parB Caulobacter* cells **(A)** without an extra ectopic *parS_3+4_*, **(B)** with an extra ectopic *parS3+4* at +200 kb, **(C)** at +1000 kb, or **(D)** at +1800 kb. Cell length of an exponentially-growing cells were quantified and presented as histograms. Vertical dotted red lines indicate the mean cell length. The number of CFP-ParB foci (green) per cell was also quantified and plotted as histograms. Note that we could not observe foci corresponding to an extra ectopic *parS3+4* perhaps due to the limited numbers of ParB bound to this shorter cluster. Most observable foci are likely due to the original *parS1-7* cluster that reside ~8 kb near ori.

### Systematic identification of a permissive zone for *parS* insertion by transposon mutagenesis with deep sequencing (Tn-seq)

Previously, a comparative genomics study surveyed and predicted the positions of *parS* sites over a wide range of bacteria and found that most *parS* sites are located close to the *ori* on the chromosome (7). Here, in *Caulobacter*, we have found that a second *parS* cluster, depending on its location on the chromosome, can affect chromosome segregation and cell fitness. To investigate this positional bias systematically, we employed a genome-wide transposon mutagenesis with deep sequencing (Tn-seq) approach. Briefly, a Tn5 transposon carrying *parS* sites 3, 4 and 5 was used to insert these strong *parS* sites randomly around the chromosome. A library of approximately half a million of single colonies were generated and the genomic locations of the inserted *parS* cluster was then determined *en masse* by deep sequencing. As a control, we generated an insertion library using a transposon that does not carry *parS*. Wild-type *Caulobacter* cells were first mutagenized with *parS*^+^ or *parS*^−^ transposon, and the number of insertions was binned to 10-kb segments along the *Caulobacter* chromosome. The ratio of the frequency for the *parS*^+^ transposon and that of the *parS*^−^ transposon was plotted as a log_10_ scale against genomic position (Fig. 7A), and used as a proxy to determine the genomic preference for an extra cluster of *parS*. We observed that a second *parS* cluster is most tolerated within ~500 kb surrounding *ori* (Fig. 7A and Fig. S8A). In contrast, an ectopic *parS* is strongly disfavoured near the middle of each chromosomal arm (Fig. 7A and Fig. S8B), consistent with our observation that *parS*^3+4^ at +1000 kb caused cell elongation and chromosome segregation defects. A limited zone of *parS* enrichment was also found within ~100 kb around the *ter* (Fig. 7A and Fig. S8C). Lastly, we also note the presence of two *parS* insertion “hot spots”. The first hot spot locates near the native *parS* cluster (Fig. 7B), likely strengthening the existing native ParB binding area on the left arm of the chromosome. The second hot spot encompasses the *recF*, *gyrB* and *CCNA0160* genes (Fig. 7C). One possibility is that a *parS* insertion in the vicinity of *gyrB* is preferred because it alters the global supercoiling level. However, we found that the *gyrB* transcription was unchanged compared to wild-type cells or cells with an extra *parS* elsewhere on the chromosome. The mechanism responsible for the *gyrB* “hotspot” therefore remains unknown.

**Figure 7.**
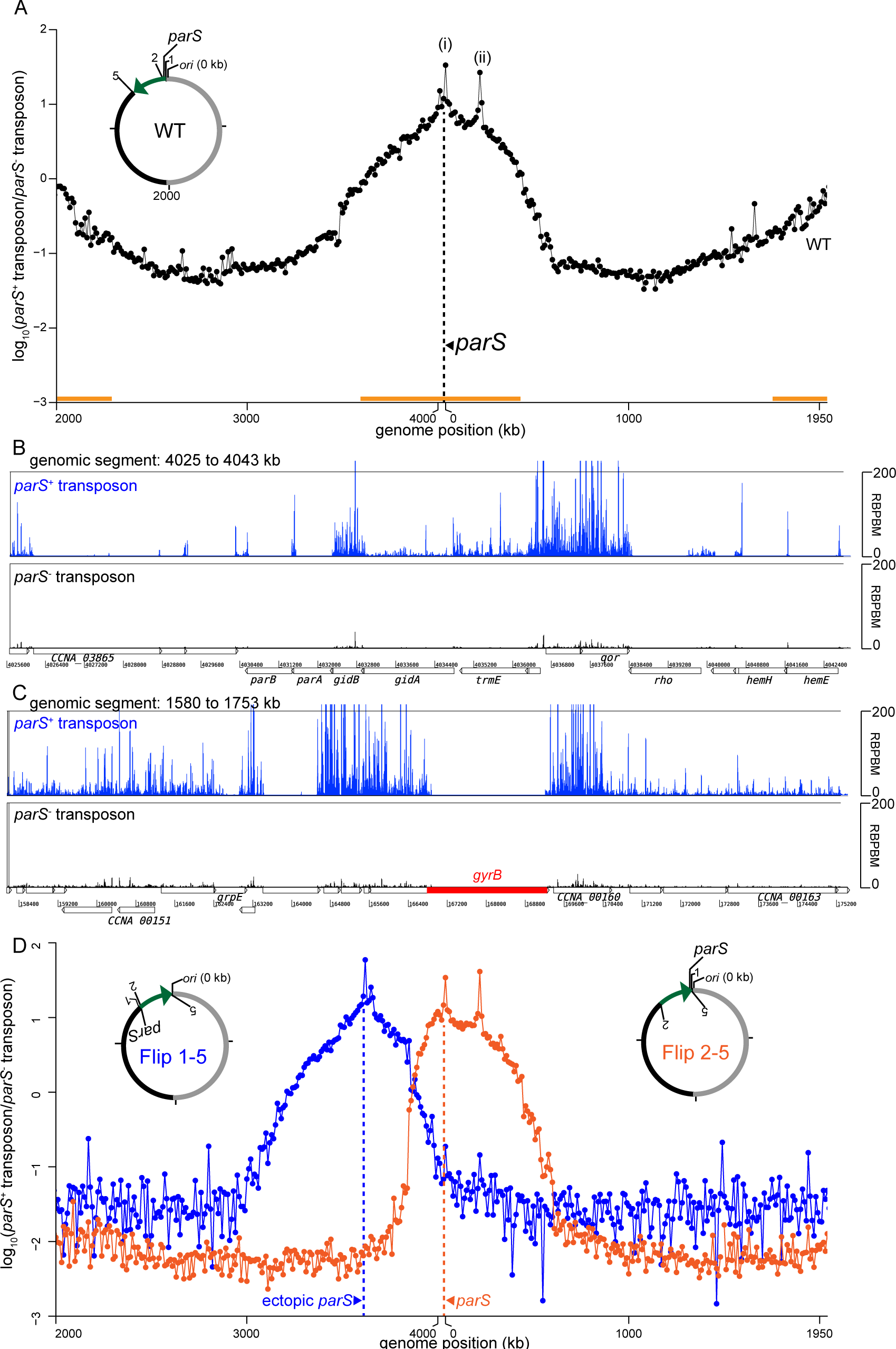
Tn5-seq reveals the positional bias of the centromeric *parS* site on *Caulobacter* chromosome. **(A)** Wild-type *Caulobacter* cells were mutagenized with the *parS+* or *parS-* transposon, and the number of insertions was binned to 10-kb segments along the genome. The ratio between insertion frequency for the *parS+* transposon and that of the *parS-* transposon was calculated and plotted as a log10 scale against genomic position. Two hotspots for insertion of the *parS+* transposon are marked with asterisks (*). The vertical dotted line (black) shows the position of the native *parS* cluster. The horizontal bar (orange) indicates the permissive zone for extra *parS* insertions. **(B-C)** Comparison between *parS+* (blue) and *parS-* (black) transposon insertions for the genomic segment between +4025 kb and +4043 kb, and between +158 kb and +175 kb. **(D)** *parS+*/*parS-* Tn5-seq profiles for Flip1-5 (blue) and Flip 2-5 (orange) strains. The horizontal axis represents genome position in kilobases for each strain. A schematic genomic map of Caulobacter showing the position of *parS* and *ori* are presented in the inset. The inverted DNA segment (green arrow) is indicated together with the end points of the inversion (1, 2, and 5).

We noted that *parS* insertion frequency decreases gradually from *ori* to the mid-arm without a clear boundary, suggesting that the *parS* permissive zone is perhaps dependent on the genomic distance away either from *ori* or from the native *parS* cluster. To test this hypothesis, we employed a Flip 1-5 strain where the native cluster of *parS* sites were relocated ~400 kb away from *ori* through an inversion between +3611 kb and +4038 kb (Fig. 7D) (39). The Tn5 transposon with or without the *parS* cluster was again used to randomly mutagenize the Flip 1-5 strain. As a control, we also transposon mutagenized another inversion strain (Flip 2-5) where the native *parS* cluster remains at its original location but a similar chromosome segment (between +3611 kb and +4030 kb) was inverted (Fig. 7D). Results showed that the permissive zone for insertion of an extra *parS* cluster in Flip 1-5 was now centred near the relocated *parS* site at +3611 kb, while the permissive zone remains centred at the native *parS* in the control Flip 2-5 strain (Fig. 7D) (39). Altogether, our results suggest that the genomic distance from the original *parS* cluster, not the distance from *ori*, is likely the main determinant of the permissive zone for the insertion of a second *parS* cluster.

Most bacterial species with a ParAB-*parS* system have more than one *parS* site (7), and some species such as *Streptomyces coelicolor* and *Listeria innocua* have accumulated 22 *parS* sites near their origin of replication (7, 40). How the bacterial centromere-like region expands and what drives its extension over time are interesting biological questions. Our finding that new *parS* sites can locate near the native *parS* cluster but not elsewhere could potentially explain the clustering of *parS* sites on bacterial chromosomes over time. New *parS* sites preferentially locate near the original *parS* cluster because it is the least disruptive to chromosome segregation, cell division, and cell viability (Fig. 6 and 7). In *Caulobacter*, *parS,* not *ori*, is the site at which force is exerted during chromosome segregation (8). ParA forms a gradient emanating from the opposite pole to the ParB-*parS* cluster. A ParA gradient retracts upon contacting ParB-*parS* and this nucleoprotein complex moves in the retreating gradient of ParA to the opposite cell pole. ParA-ParB-*parS* are only required for the segregation of *parS*-proximal DNA, but not of the distal DNA loci (41). Once the *parS*-proximal DNA is properly segregated by ParA-ParB-*parS*, distal DNA regions follow suit, driven by separate molecular machinery, or more likely without the need of a dedicated system (41). It is, therefore, foreseeable that expanding the *parS* region by adding new *parS* sites near the native cluster is least disruptive to chromosome segregation and the subsequent cell division since the *parS*-proximal DNA remains the first locus to be segregated. Similarly, in *P. aeruginosa*, *parS* is also the first segregated locus and it is preferable for cell viability that *parS* segregates soon after DNA replication (5).

In this study, we also discovered that new *parS* sites are also tolerated near the *ter* region, albeit with less preference than near the native *parS* cluster. In *P. aeruginosa* or *B. subtilis*, insertion of *parS* near the *ter* region is strongly discouraged, presumably due to the recruitment of the Structural Maintenance of the Chromosomes (SMC) complex away from *ori* (5, 42). SMC is a prominent protein involved in bacterial chromosome organization and segregation (39, 42–45). To test if SMC might contribute to shape the distribution of ectopic *parS* sites in *Caulobacter*, we transposon mutagenized the *Δsmc Caulobacter* strain (Fig. S8D). In *Δsmc* cells, the pattern of *parS* permissive zones does not change dramatically. New *parS* sites remain disfavoured near mid-arms, although they are less favoured near *ter* compared to wild-type cells (Fig. S8D). Our previous study showed that *Caulobacter* SMC are recruited to the *ter*-located ectopic *parS* and cohese flanking DNA together, nevertheless the global chromosome organization remained largely unchanged with *ori* and *ter* at opposite poles and two chromosomal arms running in parallel down the long axis of the cell (39). All together, we conclude that SMC contributes to the determination of *parS* permissive zones but cannot solely explain some of the preference for the *ter* region and the disfavour for mid-arm regions in *Caulobacter crescentus*. Further investigation into the molecular mechanism that gives raise to the permissive zones of *parS* will undoubtedly improve our understanding of bacterial chromosome segregation and organization.

## SUPPLEMENTARY DATA

Supplementary Data are available online.

## ACCESSION NUMBERS

The accession number for the sequencing data reported in this paper is GEO: **GSE100233**.

## ACKNOWLEDGMENTS

We thank Anjana Badrinarayanan, Matthew Bush, Mark Buttner, Michael Laub, and Susan Schlimpert for discussion and comments on the manuscript. We thank Christine Jacob-Wagner and Martin Thanbichler for materials.

## FUNDINGS

This study was supported by a Royal Society University Research Fellowship (UF140053), a Royal Society Research Grant (RG150448), a BBSRC Grant (BB/P018165/1) to T.B.K.L. and a BBSRC grant-in-add (BBS/E/J/000C0683) to the John Innes Centre.

### Conflict of interest statement

None declared.

